# Plasma neurofilament light protein provides evidence of accelerated brain ageing in treatment-resistant schizophrenia

**DOI:** 10.1101/2023.11.06.565715

**Authors:** Cassandra M. J. Wannan, Dhamidhu Eratne, Alexander F. Santillo, Charles Malpas, Brandon Cilia, Olivia M. Dean, Adam Walker, Michael Berk, Chad Bousman, Ian Everall, Dennis Velakoulis, Christos Pantelis, The MiND Study Group

**Affiliations:** Melbourne Neuropsychiatry Centre, Department of Psychiatry, The University of Melbourne, Melbourne, VIC, Australia; Centre for Youth Mental Health, The University of Melbourne, Parkville, Victoria, Australia; Orygen, Parkville, Victoria, Australia; Neuropsychiatry, The Royal Melbourne Hospital, Melbourne, VIC, Australia; Department of Clinical Sciences, Clinical Memory Research Unit, Faculty of Medicine, Lund University, Lund/Malmö, Sweden; Department of Medicine, Royal Melbourne Hospital, University of Melbourne; Melbourne School of Psychological Sciences, University of Melbourne; Deakin University, IMPACT – the Institute for Mental and Physical Health and Clinical Translation, School of Medicine, Barwon Health, Geelong, Australia; Florey Institute of Neuroscience and Mental Health, Parkville, Australia; Department of Medical Genetics, University of Calgary, Calgary, Canada; Department of Psychiatry, University of Melbourne, Melbourne, Australia; Institute of Psychiatry, Psychology & Neuroscience, King’s College London, London, UK; NorthWestern Mental Health, Sunshine Hospital, Melbourne, VIC, Australia

## Abstract

**Background:** Accelerated brain aging has been observed across multiple psychiatric disorders. Blood markers of neuronal injury such as Neurofilament Light (NfL) protein may therefore represent biomarkers of accelerated brain aging in these disorders. The current study aimed to examine whether relationships between age and plasma NfL were increased in individuals with primary psychiatric disorders compared to healthy individuals.

**Methods:** Plasma NfL was analysed in major depressive disorder (MDD, n = 42), bipolar affective disorder (BPAD, n = 121), treatment-resistant schizophrenia (TRS, n = 82), a large reference normative healthy control (HC) group (n= 1,926) and a locally-acquired HC sample (n = 59). A general linear model (GLM) was used to examine diagnosis by age interactions on NfL z-scores using the large normative HC sample as a reference group. Significant results were then validated using the locally-acquired HC sample.

**Results:** a GLM identified a significant age by diagnosis interaction for TRS vs HCs and BPAD vs HCs. Post hoc analyses revealed a positive correlation between NfL levels and age among individuals with TRS, whereas a negative correlation was found among individuals with BPAD. However, only the TRS findings were replicated using the locally-acquired HC sample. Post hoc analyses revealed that individuals with TRS aged <40 had lower NfL levels compared to same-age HCs, whereas individuals with TRS aged >40 had higher NfL levels compared to same-age HCs.

**Conclusions:** These findings add to the growing literature supporting the notion of accelerated brain ageing in schizophrenia-spectrum disorders.

## Introduction

Structural and functional brain abnormalities have consistently been observed in psychiatric disorders such as depression, anxiety, bipolar disorder, and schizophrenia^1,2^. Recent studies indicate that subtle neuroprogressive brain changes occur across the course of these illnesses, with one possible explanation for these findings being accelerated ageing^3^.

Increased brain age gaps, a marker of accelerated brain ageing, have been observed in individuals with schizophrenia, major depression, borderline personality disorder, bipolar disorder, and in those at ultra high risk for psychosis^3,4^. Across these studies, accelerated brain ageing was greatest in individuals with schizophrenia. These cross-sectional findings are consistent with a longitudinal study which found that individuals with schizophrenia not only had increased brain age at baseline, but also that the brain age gap increased over a 4-year follow-up period ^5^. Together, these findings support the notion that individuals with serious mental illness experience accelerated brain ageing, with the greatest impact seen in those with schizophrenia.

Given evidence for accelerated brain ageing in psychiatric disorders, numerous studies have sought to determine the potential mechanisms and consequent biomarkers underlying this process. One possible biomarker is neurofilament light (NfL). NfL forms an essential component of the neuronal cytoskeleton particularly important for growth and stability of axona^6^. It also is a marker of potential neuronal injury. While some studies have observed increased NfL in schizophrenia and depression^7,8^, recent studies from our group using ultrasensitive technology to measure plasma NfL levels found no evidence of increased NfL in treatment-resistant schizophrenia (TRS) or major depressive disorder (MDD), but a slight increase in individuals with bipolar affective disorder (BPAD)^9,10^. However, these studies did not examine whether age trajectories of NfL in psychiatric disorders differed from those seen in healthy individuals, where higher NfL levels are associated with increasing age ^11^. Given findings of accelerated brain ageing in schizophrenia, BPAD, and MDD, it is possible that NfL levels would increase at a faster rate in individuals with psychiatric disorders compared to healthy individuals. If this is the case, NfL could potentially represent a theragnostic, stage, or prognostic marker^12^ in psychiatry where there is a dearth of valid single marker assays. The aim of the current study was therefore to examine whether relationships between age and plasma NfL were increased in individuals with primary psychiatric disorders compared to healthy individuals.

## Methods

### Participants

Participant samples and data, described in detail previously ^9,13–16^, were included from three patient cohorts and two control groups. Patient cohorts included individuals with BPAD (n = 121), MDD; n = 42, and TRS, n = 82. Healthy control (HC) groups included a large reference normative control group (n= 1,926) and a local sample (n = 59) with no current or past psychiatric or neurological illness.

### Sample analysis

Plasma aliquots from all samples were stored at −80°C. Samples were randomised before analysis, and analyses blinded to diagnosis. Plasma NfL levels were measured on Quanterix SR-X and HD-X analysers using Simoa NF-Light kits, according to the manufacturer’s recommendations.

### Data analysis

Data were analysed using R version 4.2.2.

#### Calculation of z-scores

NfL Z-scores were calculated from age-adjusted percentiles from the large reference cohort (Control Group 2), derived using generalised additive models for location, scale, and shape ^9^.

#### Examination of the impact of diagnosis on relationships between age and NfL

Primary analyses were conducted using the large reference normative control group. A general linear model (GLM) was used to examine diagnosis by age interactions on NfL z-scores, with HCs as the reference group and sex added as a covariate. Where significant diagnosis by age interactions were observed, post-hoc analyses were performed to examine relationships between age and NfL levels separately in the relevant diagnostic groups. We also examined relationships between NfL levels and polynomial contrasts for age (up to the fourth order) to examine non-linear relationships, however, relationships were strongest and Bayesian information criteria lowest for raw age values and we therefore present these findings.

For all analyses, 95% confidence intervals were computed (nonparametric bootstrapping, 1000 replicates), with statistical significance defined as any confidence interval not including the null (at 95% level).

Replication analyses were conducted using the local healthy control group in order to (1) examine the robustness of our findings using a distinct HC sample, and (2) determine the impact of body weight, which was not available in the large reference group, on the relationship between age and NfL, and diagnosis by age interactions.

#### Post-hoc analyses

In post-hoc analyses, for diagnoses where significant significant diagnosis by age interactions were observed, age grouping was performed based on the mean age for the relevant diagnostic group. GLMs were then used to examine between-group differences in NfL z-scores in diagnostic groups vs healthy controls separately in older and younger individuals, adjusting for sex.

In addition, relationships between NfL z-scores and and in TRS individuals were further interrogated while adjusting for clozapine dose, chlorpromazine equivalent dose, and serum clozapine levels, adjusting for age and sex, in order to determine whether antipsychotic medication impacted on relationships between age and NfL levels.

## Results

### Examination of the impact of diagnosis on relationships between age and NfL

As seen in Figure 1, a GLM identified a significant age by diagnosis interaction for TRS vs HCs (*β = 0*.*05 [0*.*01, 0*.*08], p* < .001) and BPAD vs HCs (*β = -0*.*02 [-0*.*04, -0*.*005], p* = .007). Post hoc analyses revealed a positive correlation between NfL levels and age among individuals with TRS (*β = 0*.*05 [0*.*02, 0*.*09], p* = .005), reflective of an accelerated increase. Conversely, a negative correlation was found among individuals with BPAD (*β = -0*.*02 [-0*.*04, -0*.*01], p* = .009), indicative of a slower increase in NfL levels with increasing age.

**Figure 1.**
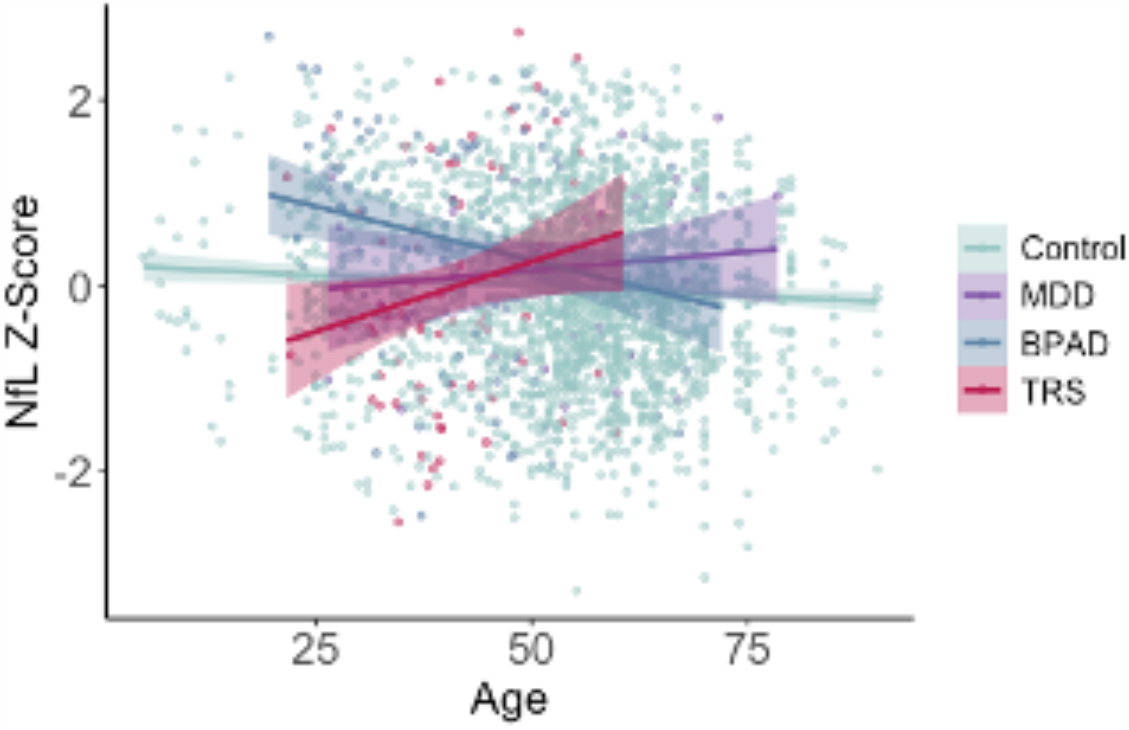
Relationships between age and age-normalized NfL z-scores

**Figure 2.**
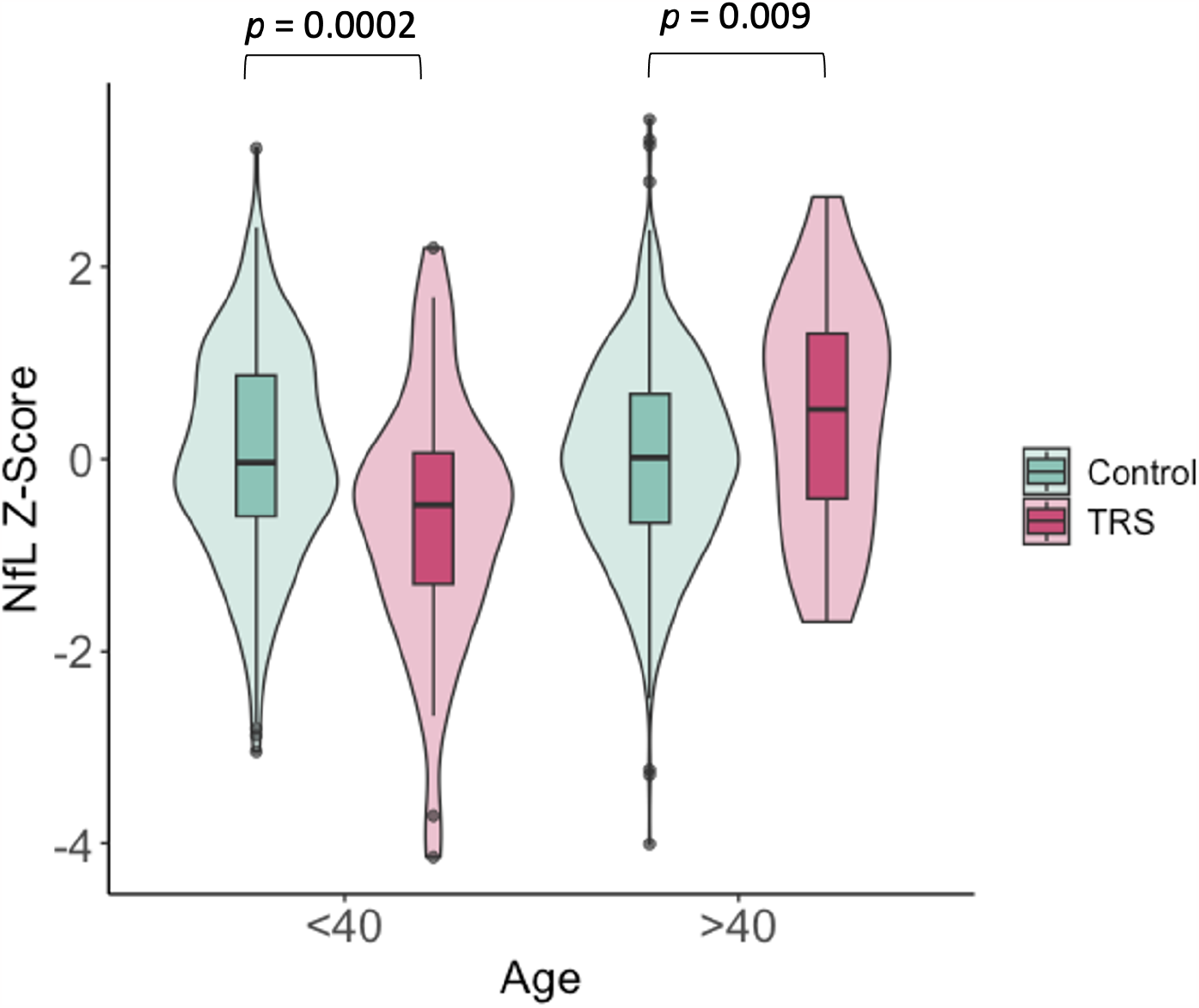
Comparisons of neurofilament light protein (NfL) levels in TRS individuals separated by age. TRS individuals aged <40 had significantly lower NfL levels compared to a large reference cohort of healthy controls, whereas TRS individuals aged >40 had higher NfL levels compared to healthy controls.

In secondary analyses using a sample of locally-acquired healthy controls, the diagnosis x age interaction was still present for the TRS cohort relative to HCs (*β = 0*.*06 [0*.*03, 0*.*11], p* = .002). However, the diagnosis by age interaction was not significant for BPAD vs HCs (*β = 0*.*002 [0*.*03, -0*.*02], p* = .899). The diagnosis by age interaction for TRS individuals vs locally-acquired healthy controls also remained significant when controlling for weight (*β = 0*.*06 [0*.*03, 0*.*11], p* = .002)

### Supplementary analyses

In supplementary analyses, younger individuals with TRS (age < 40) had significantly lower NfL levels than HCs (*β = -0*.*56 [-0*.*98, -0*.*21], p* < .001). Conversely, older individuals (age > 40) with TRS had significantly higher NfL levels than older HCs (*β = 0*.*49 [0*.*08, 0*.*92], p* = .001). Younger individuals with BPAD had significantly higher NfL levels than HCs (*β = 0*.*56 [0*.*27, 0*.*87], p* < .001) whereas NfL levels in older individuals with BPAD did not differ from HCs (*β = 0*.*28 [-0*.*01, 0*.*59], p* = .04).

The positive relationships between age and NfL z-scores in individuals with TRS remained significant when controlling for clozapine dose (*β = 0*.*04 [0*.*01, 0*.*8], p* = .01), chlorpromazine equivalent dose (*β = 0*.*04 [0*.*01, 0*.*8], p* = .009), and serum clozapine levels (*β = 0*.*04 [0*.*01, 0*.*8], p* = .01).

## Discussion

Compared to a large normative sample of healthy individuals, individuals with TRS demonstrated a stronger positive relationship between age and age-normalized NfL levels, whereas individuals with bipolar disorder demonstrated a stronger negative relationship. The TRS findings were replicated when using a locally-acquired healthy control sample, however, the attenuated age-NfL relationship in individuals with BPAD was not replicated in this secondary analysis.

Stronger relationships between age and NfL levels in individuals with TRS may provide further evidence for accelerated brain ageing in individuals with schizophrenia-spectrum disorders, and specifically those with treatment-resistant illness. Importantly, after inclusion of clozapine dose, chlorpromazine equivalent dose, and serum clozapine levels as covariates in these analyses, relationships between age and age-normalized NfL levels remained significant, indicating that these relationships are not driven by antipsychotic medication. While previous neuroimaging studies have provided consistent evidence for accelerated brain ageing in schizophrenia^3–5,17^, other biomarker studies have provided less consistent findings^18^. Thus, confirmation of our findings in larger samples, including those with treatment-responsive schizophrenia, will be critical to confirm this accelerated age-related increase in NfL in TRS.

A limitation of the current study is its cross-sectional design: although we can posit that stronger relationships between age and NfL is indicative of accelerated brain ageing in TRS, further longitudinal studies are needed to fully interrogate this hypothesis. Additionally, NfL values from our large normative healthy sample were obtained from a different laboratory than our psychiatric and locally-acquired healthy control groups, which may lead to site-related differences. However, validation of our TRS finding in our locally-acquired HCs suggests that these findings are robust to any potential site differences.

In summary, this study found that the relationship between age and NfL levels was stronger in individuals with TRS compared to two independent samples of healthy individuals. These findings may add to the growing literature supporting the notion of accelerated brain ageing as part of the process of neuroprogression in schizophrenia-spectrum disorders^19^.

## Acknowledgements

The author(s) disclosed receipt of the following financial support for the research, authorship, and/or publication of this article: The authors acknowledge the financial support of the CRC for Mental Health. The Cooperative Research Centre (CRC) programme is an Australian Government Initiative. The authors acknowledge the CRC Scientific Advisory Committee, in addition to the contributions of study participants, clinicians at recruitment services, staff at the Murdoch Children’s Research Institute, staff at the Australian Imaging, Biomarkers and Lifestyle Flagship Study of Aging, and research staff at the Melbourne Neuropsychiatry Centre, including coordinators Merritt, A., Phassouliotis, C., and research assistants, Burnside, A., Cross, H., Gale, S., and Tahtalian, S. Participants for this study were sourced, in part, through the Australian Schizophrenia Research Bank (ASRB), which is supported by the National Health and Medical Research Council of Australia (Enabling Grant N. 386500), the Pratt Foundation, Ramsay Health Care, the Viertel Charitable Foundation and the Schizophrenia Research Institute. We thank the Chief Investigators and ASRB Manager: Carr, V., Schall, U., Scott, R., Jablensky, A., Mowry, B., Michie, P., Catts, S., Henskens, F., Pantelis, C., Loughland, C. We acknowledge the help of Jason Bridge for ASRB database queries. The authors are grateful for assistance from Brett Trounson and Dr Christopher Fowler and the team at The Florey Oak St Biobank.

MB is supported by a NHMRC Senior Principal Research Fellowship and Leadership 3 Investigator grant (1156072 and 2017131). A.J.W. was supported by a Trisno Family Fellowship, funded in part by an NHMRC CRE (1153607). C.P. was supported by a National Health and Medical Research Council (NHMRC) Senior Principal Research Fellowship (1105825), an NHMRC L3 Investigator Grant (1196508). This study was also supported by MACH MRFF RART 2.2, NHMRC (1185180), and Psychiatry and Rehabilitation Division, Region Sk.ne, Sweden. The role of these funding sources was to support research study staff and biosample analyses.

